# Sublethal immune resistance to parasites generates reaction-norm patterns indistinguishable from tolerance

**DOI:** 10.64898/2026.06.30.735575

**Authors:** Otto Seppälä, Ben Ashby

## Abstract

Hosts defend themselves against parasites through resistance (reducing parasite burden) and tolerance (reducing the fitness cost of infection without affecting parasites). This distinction has important evolutionary implications: resistance is predicted to maintain polymorphism while tolerance tends to fix, and only resistance is expected to provoke parasite counter-adaptation. The reaction-norm framework, which infers tolerance from the slope of host fitness regressed on parasite burden, assumes that a shallow slope reflects parasite-independent host protection. We test this assumption using a within-host model in two variants: microparasites (Model 1, with within-host replication) and macroparasites (Model 2, without). Sublethal immunity impairs the parasite’s host-exploitation rate, reducing both growth and per-parasite virulence without killing them. We show that this generates systematic slope differences among host genotypes that the framework interprets as variation in tolerance. Furthermore, the ranking of genotypes’ slopes reverses between linear and sigmoidal damage functions: under linear damage, the strongest immune responder appears most tolerant; under sigmoidal damage, the weakest responder does. Decomposition of the damage reduction shows that virulence reduction accounts for the majority of the effect across both model variants. Thus, the reaction-norm slope cannot determine whether host fitness is maintained by parasite-independent tissue protection or by sublethal impairment of parasites.

## Introduction

Parasites reduce host fitness, and hosts have evolved two fundamentally different strategies to counter them. Resistance reduces parasite burden through immune killing, while tolerance reduces the fitness cost of infection without affecting parasite numbers (Schneider & Ayres, 2008; Råberg et al., 2009). The key distinction is that tolerance is parasite-independent: the host limits damage to its own tissues without harming the parasite. This dichotomy was first observed in plant pathology by Cobb (1894), who noted that some wheat varieties produced acceptable yields despite heavy rust infection, and was defined as the resistance–tolerance distinction by Schafer (1971). Råberg et al. (2007) extended the framework to animals, and it has since become a cornerstone of host–parasite evolutionary ecology, shaping how researchers measure defence, model its evolution, and predict epidemiological consequences.

The distinction carries important theoretical implications. Roy and Kirchner (2000) showed that resistance and tolerance have contrasting evolutionary dynamics. Resistance creates negative frequency-dependent selection and maintains polymorphism, whereas tolerance tends to fix in populations. Furthermore, because tolerance, by definition, does not reduce parasite fitness, it is predicted to increase disease prevalence (Boots, 2008; Miller et al., 2006). The coevolutionary implications are equally divergent. Resistance is expected to provoke counter-adaptation from parasites, whereas tolerance, because it does not directly affect parasite fitness, is not (Råberg et al., 2009).

Current research on tolerance rests heavily on the reaction norm framework, which measures tolerance as the slope of host fitness regressed on parasite burden, a shallow slope indicating that the host maintains fitness despite high parasite load (Råberg et al., 2007; Simms, 2000). The framework assumes that a shallow slope reflects host processes that do not affect the parasite. But immune responses are not binary. Between the extremes of no response and killing lies a broad range of intensities at which the immune system damages parasites without eliminating them. A host operating in this range would reduce per-parasite harm and, depending on whether the parasite reproduces within the host, may also modestly reduce parasite density. Such effects could potentially appear as tolerance within the reaction norm framework, despite relying on parasite-directed defence.

Several lines of evidence suggest this may be common. Ayres and Schneider (2008) showed that a single mutation in *Drosophila* alters both resistance and tolerance as measured by the reaction norm, and that the same physiological response—anorexia—has opposite effects depending on the pathogen, increasing tolerance to one infection while decreasing resistance to another (Ayres & Schneider, 2009). Louie et al. (2016) showed that realistic tolerance curves are nonlinear, requiring multiple parameters rather than a single regression slope, and Torres et al. (2016) argued that single-time-point regressions miss the multi-dimensional trajectories of infection dynamics. These observations raise a question that has been discussed qualitatively (Ayres & Schneider, 2012; Kutzer & Armitage, 2016) but, to our knowledge, not formally modelled: can the reaction-norm framework identify whether a shallow slope reflects reduced parasite density, reduced per-parasite virulence, or both?

We address this question using a mathematical model built around a single empirical observation: many immune effectors damage parasites before killing them. For example, reactive oxygen species produced by activated macrophages are bactericidal at high concentrations, but at lower concentrations they impair specific metabolic pathways by oxidising iron–sulfur cluster enzymes, slowing growth without killing (Fang, 2011; Jang & Imlay, 2007). Hepcidin-mediated iron sequestration restricts microbial access to iron, slowing pathogen replication without directly killing, and—for virulence factors that require iron as a cofactor—reducing their activity (Drakesmith & Prentice, 2012; Ganz & Nemeth, 2015; Palmer & Skaar, 2016). Furthermore, antibodies against merozoite surface proteins can both opsonise parasites for phagocytic destruction and inhibit erythrocyte invasion, with the relative contribution of these mechanisms depending on antibody specificity and IgG subclass (Beeson et al., 2016; Teo et al., 2016). In each case, the same effector molecule produces qualitatively different outcomes depending on its concentration: killing above a threshold, impairment below it. In the model, we capture this with two dose-response functions: sublethal impairment of parasite condition begins at any level of immune activity, whereas killing requires activity above a critical intensity. We show that sublethal impairment alone generates reaction-norm patterns indistinguishable from tolerance.

## Model

We develop a within-host model in which immunity impairs parasites sublethally before killing begins. Both parasite growth rate and virulence decline as downstream consequences of this impairment, but the two traits need not respond with equal sensitivity. The model, therefore, uses separate impairment parameters *β_r_* and *β_α_* for growth and virulence, respectively. We analyse the model in two variants: microparasites (Model 1, with within-host reproduction) and macroparasites (Model 2, no within-host reproduction).

### Immune effector functions

Consider a host with immune intensity *i* ∈ [0, 1] against a parasite infrapopulation of density *P*. The variable *i* is an abstract, dimensionless quantity representing the magnitude of immune capacity, normalised so that *i* = 0 represents no immune activity and *i* = 1 represents maximal defence. Real immune defence is multidimensional, but we collapse this complexity into a scalar for tractability. The immune response can impair two aspects of parasite performance—growth rate and virulence—with potentially different sensitivities:

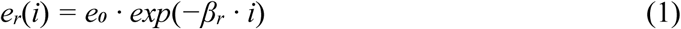

and

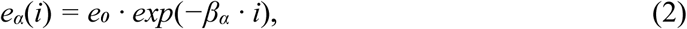

where *e₀* is the baseline exploitation rate, and *β_r_* and *β_α_* are the impairment sensitivities for growth and virulence, respectively. The effective growth rate and virulence are downstream consequences of exploitation:

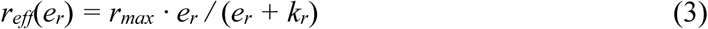

and

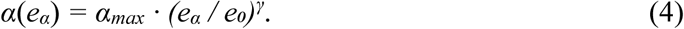

The growth rate *r_eff_* follows a saturating (Michaelis-Menten) function of the growth-exploitation component: at high *e_r_*, growth approaches *r*_max_; as *e_r_* declines, growth rate falls. The parameter *k_r_* is the half-saturation constant. Virulence *α* follows a power law: per-parasite harm scales with the *γ*-th power of relative exploitation. When *γ* = 1, virulence is proportional to exploitation. When *γ* > 1, virulence is a disproportionately accelerating function, because the most harmful aspects of exploitation (e.g., tissue destruction, toxin production) require a high exploitation rate. When *γ* < 1, virulence is a decelerating function. We use *γ* = 1 as the default.

This framework contains three special cases: (1) *β_r_* = *β_α_*: immunity targets a single exploitation rate from which both growth and virulence decline together; (2) *β_r_* > 0, *β_α_* = 0: immunity reduces growth without reducing per-parasite virulence; (3) *β_r_* = 0, *β_α_* > 0: immunity reduces virulence without affecting growth or density. Intermediate combinations represent mechanisms that affect both traits to different degrees.

The immune killing rate follows a Hill function with a steep threshold:

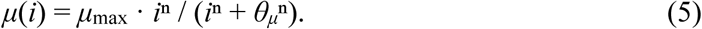

Here *μ*_max_ is the maximum killing rate, *θ_μ_* is the half-maximal activation threshold, and *n* is the Hill coefficient controlling steepness. Killing remains negligible at low *i* and rises sharply near the threshold. This function is independent of the impairment parameters and applies identically in both model variants.

### Within-host dynamics

The two model variants differ in whether the parasite reproduces within the host (Model 1, microparasites) or establishes from the environment without within-host reproduction (Model 2, macroparasites).

Model 1: within-host dynamics of a microparasite (with within-host replication) follow logistic growth with immune-mediated impairment and killing:

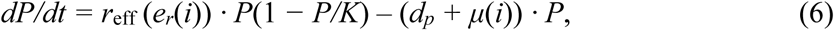

where *K* is the within-host carrying capacity (the maximum parasite density in the absence of immune defence, determined by resource availability and spatial constraints) and *d_p_* is the baseline parasite mortality rate. The equilibrium parasite density is:

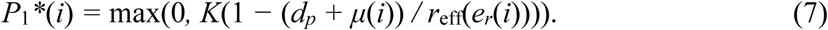

This equilibrium is positive whenever *d_p_* + *μ*(*i*) < *r*_eff_(*e_r_*(*i*)). When *d_p_* + *μ*(*i*) ≥ *r*_eff_(*e_r_*(*i*)), the parasite cannot replace its losses and the infection is cleared (*P*\* = 0). Note that sublethal impairment reduces *P*_1_* even without killing (*μ* = 0) because parasites die at a rate *d_p_* regardless of immune activity, and a more slowly-growing parasite cannot replace these losses as efficiently. The magnitude of this effect depends on the ratio *d_p_*/*r*_eff_(*e_r_*(*i*)).

<u>Model 2</u>: within-host dynamics of a macroparasite (no within-host replication) is determined by the establishment of parasites from the environment at rate Λ (produced descendants are released to the environment as transmission stages). The establishment of new parasites is opposed by the same immune effectors that act on established ones (Brunet et al., 1999; Smithers & Terry, 1969; Anthony et al., 2007). We therefore model the establishment rate as Λ(*i*) = Λ₀ · (*r*_max_/*r*_max,ref_) · (1 − *μ*(*i*)/*μ*_max_), where Λ₀ is the establishment rate in the absence of immune defence. The establishment rate is clone-specific, scaling with each clone’s maximum growth rate *r*_max_ (Eq. 3) relative to the reference (medium) clone *r*_max_,_ref_ = 1.20. Because within-host density is independent of growth in Model 2, this fecundity scaling plays the role that the growth function (Eq. 7) plays for the clones in Model 1. Within-host dynamics are:

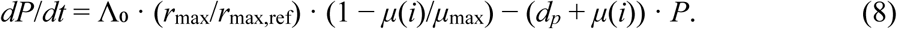

The equilibrium parasite density is:

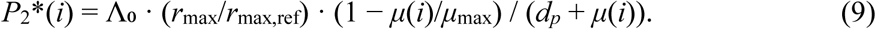

Because both the establishment term and the mortality term depend on *μ*(*i*), which is small below its activation threshold *θμ*, the sublethal resistance zone has *P*_2_* ≈ Λ₀(*r*_max_/*r*_max,ref_)/*d_p_*. The host’s immune activity degrades the parasites it harbours without reducing their number. Only when immune intensity crosses the killing threshold does *P*_2_* decline, through a simultaneous reduction in establishment and an increase in mortality of the established parasites. Sublethal impairment affects only each parasite’s condition, with implications for virulence, but not parasite density.

### Host fitness

Host fitness is determined by the balance between parasite-induced damage and the cost of immune activity:

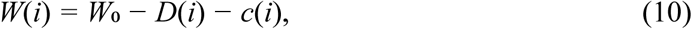

where *W*₀ is the baseline fitness in the absence of both infection and immune expression, *D*(*i*) is the total parasite-induced damage, and *c*(*i*) is the cost of the immune activity.

#### Linear damage

Total parasite-induced damage is the product of per-parasite virulence and equilibrium parasite density:

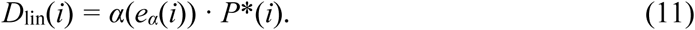

This form assumes that each parasite contributes independently and additively to host damage.

#### Sigmoidal damage

Alternatively, host damage may follow a threshold response:

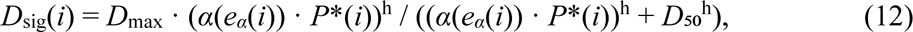

where *D*_max_ is the maximum damage achievable at saturation, *D*₅₀ is the effective damage load at which damage reaches half-maximum, and *h* controls the steepness of the threshold. When *h* = 1, this reduces to a Michaelis-Menten saturating response (i.e., damage increases with load but plateaus at high loads). When *h* is large, the response is switch-like: the host tolerates parasites with little fitness cost up to a threshold, beyond which fitness collapses. This form captures the biological reality that many disease outcomes are nonlinear (Louie et al., 2016).

The immune cost function is assumed to accelerate with increasing *i*:

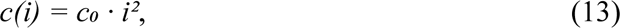

where *c*₀ is the cost coefficient. The quadratic form is a standard modelling choice representing the accelerating costs of immunopathology, autoimmune damage, and energetic investment at high intensity; accelerating defence costs of this kind select for a single intermediate level of investment (Graham et al., 2005; Boots & Haraguchi, 1999).

### Simulation analysis

We simulated an experiment in which host genotypes differ only in immune intensity *i*, with *β_r_* = *β_α_* = 2.0. This choice of *β* values is illustrative, and the full (*β_r_*, *β_α_*) parameter space is explored in the generalisation analysis. We focus the analysis on the sublethal zone of *i*, where killing is negligible, and the main immune effect is impairment of parasite condition. This provides the cleanest test of the framework’s ability to distinguish reduced per-parasite virulence from reduced parasite density. An extension to the full intensity range, including genotypes in the transition and killing zones, is presented in the Supplementary Material.

Five host genotypes were defined with mean immune intensities *ī* = 0.05, 0.10, 0.15, 0.20, and 0.25, all well below the half-maximal killing threshold *θ_μ_* = 0.6. Within-genotype variation in parasite load was generated by crossing each host genotype with three parasite clones differing in maximum growth rate: *r*_max_ = 0.72 (slow), 1.20 (medium), and 1.68 (fast). In Model 1 (microparasite), clone growth rate directly determines equilibrium density *P*\* through the growth function *r_eff_*(*e_r_*): faster clones reach higher *P*\*, although low baseline mortality keeps all clones near carrying capacity, so this separation is modest. In Model 2 (macroparasite), clones differ in fecundity, resulting in different rates of establishment from the environment (via the establishment rate Λ = Λ₀ · *r*_max_/*r*_max,ref_) and therefore different *P*\*. Each genotype-by-clone combination was replicated 20 times, with individual immune intensity drawn from a normal distribution around the genotype mean (*σ* = 0.02), with values constrained to the interval [0.01, 0.99]. Only infected individuals (*P*\* > 0.1) were included in the calculation of the reaction norm. All parameter values used in the simulations are listed in Table 1.

**Table 1.** Parameter values used throughout the analyses. Where a parameter takes one value in the main analysis and a range in the generalisation, both are given. Equation numbers refer to the Model section.

| Symbol | Description | Value |
| --- | --- | --- |
| $e_0$ | Baseline exploitation rate (Eqs 1, 2, 4) | 1.0 |
| $\beta_r$ | Growth-impairment sensitivity (Eq. 1) | 2.0 (main); 0–4 (grid) |
| $\beta_\alpha$ | Virulence-impairment sensitivity (Eq. 2) | 2.0 (main); 0–2 (grid) |
| $r_{max}$ | Maximum within-host growth rate, clone specific (Eq. 3) | 0.72, 1.20, 1.68 (slow/medium/fast clones; 1.20 used as the Model 2 normalisation reference, $r_{max,ref}$) |
| $k_r$ | Half-saturation constant for growth (Eq. 3) | 0.3 |
| $\alpha_{max}$ | Maximum per-parasite virulence (Eq. 4) | 0.12 |
| $\gamma$ | Virulence power-law exponent (Eq. 4) | 1.0 |
| $\mu_{max}$ | Maximum immune killing rate (Eq. 5) | 1.5 |
| $\theta_\mu$ | Half-maximal killing threshold (Eq. 5) | 0.6 |
| $n$ | Hill coefficient for killing (Eq. 5) | 6 |
| $K$ | Within-host carrying capacity, Model 1 (Eq. 6) | 100 |
| $d_p$ | Baseline parasite mortality rate (Eqs 6–9) | 0.05 |
| $\Lambda_0$ | Establishment rate without immunity, Model 2 (Eq. 8);<br>reference clone, clone-specific rate $\Lambda = \Lambda_0 \cdot$<br>$(r_{\max}/r_{\max,\text{ref}})$ | 5.0 (reference) |
| $W_0$ | Baseline host fitness (Eq. 10) | 100 |
| $c_0$ | Immune cost coefficient (Eq. 13) | 50 |
| $D_{\max}$ | Maximum damage, sigmoidal (Eq. 12) | 15 |
| $D_{50}$ | Half-maximal damage load, sigmoidal (Eq. 12) | 5.0 |
| $h$ | Steepness of sigmoidal damage (Eq. 12) | 4 |
| $\bar{i}$ | Genotype mean immune intensities | 0.05, 0.10, 0.15,<br>0.20, 0.25 |
| $\sigma$ | Within-genotype SD of immune intensity | 0.02 |

For each host genotype, we estimated the reaction-norm slope by regressing host fitness *W* on log₁₀(*P*\*) across all individuals of that genotype, under both linear and sigmoidal damage functions. For each genotype, we asked how much of the total reduction in parasite-induced damage came from each parasite being less harmful versus there being fewer parasites. In the reaction-norm framework, this fraction contributes to the ‘tolerance signal’. We did this using the Shapley decomposition, which averages over both possible orderings of the two channels and assigns the interaction symmetrically.

To test the generality of the results, we repeated the decomposition across a grid of (*β_r_*, *β_α_*) values ranging from 0 to 4 (*β_r_*) and 0 to 2 (*β_α_*), for both model variants and both damage functions. For each parameter combination, we recorded the fraction of the total damage reduction attributable to virulence reduction (evaluated using a single representative sublethal genotype with a mean immune intensity of 0.15 and the medium reference clone). To confirm that the slope reversal between damage functions is not specific to the choice *β_r_* = *β_α_* = 2.0, we repeated the reaction-norm analysis at four additional (*β_r_*, *β_α_*) combinations spanning different regions of the parameter space (Supplementary Figs S2–S5).

## Results

Setting *β_r_* = *β_α_* = 2.0, the reaction-norm slopes differed systematically across host genotypes in all four combinations of model variant and damage function, even though all genetic variation is in a single parasite-directed immune trait (Fig. 1). Additionally, the direction of the slope gradient reversed between the two damage functions. Under linear damage, genotypes with stronger sublethal resistance (higher *i*) had shallower slopes (Fig. 1a,c). In Model 1, the slope ranged from −22.7 (genotype with *ī* = 0.05) to −10.6 (genotype with *ī* = 0.25; Fig. 1a), which the framework would interpret as the most tolerant. Under sigmoidal damage, the ordering reversed completely (Fig. 1b), with genotypes with smaller *i* having shallower slopes (range: −7.2 to −25.1, Fig. 1b). This reversal occurs because the sigmoidal damage function places low-immunity genotypes near the saturation plateau and stronger responders on the steeper part of the curve. The same qualitative pattern held for Model 2 (macroparasite): slopes ranged from −23.3 to −12.7 under linear damage (stronger resistance → shallower slope; Fig. 1c) and from −10.2 to −26.8 under sigmoidal damage (stronger resistance → steeper slope; Fig. 1d).

**Figure 1.**
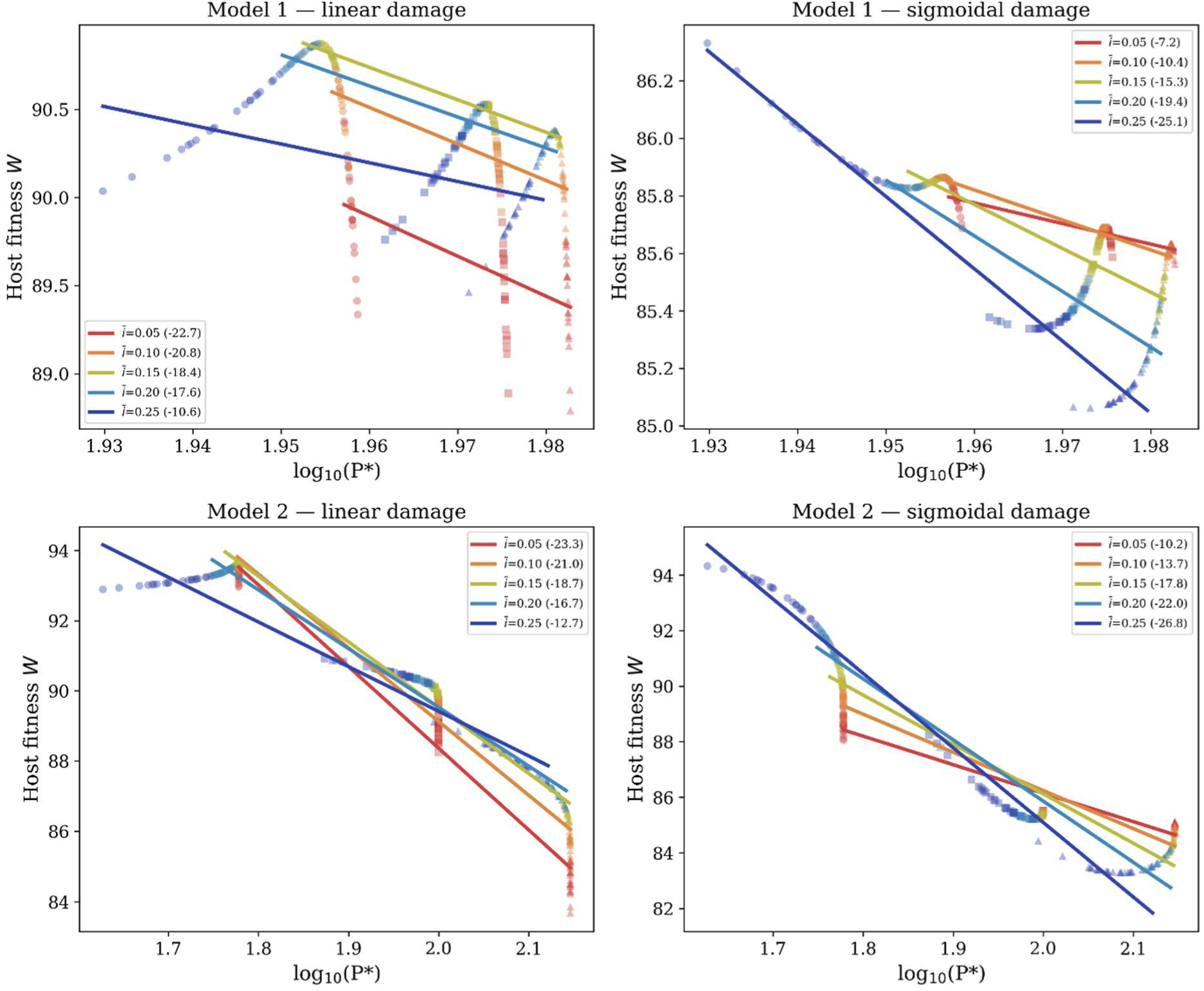
Reaction norms per host genotype for Model 1 (microparasite; top) and Model 2 (macroparasite; bottom) under linear (left) and sigmoidal (right) damage function. Markers indicate parasite clones with different reproductive rates: ○ slow, □ medium, △ fast.

Decomposition of the damage reduction of defence revealed that virulence reduction dominated in all cases, though the balance shifted between model variants (Table 2). In Model 1 (microparasite), virulence reduction accounted for 95–98% of the damage reduction of defence. In Model 2, the same range was 76–100%. The dip below 100% in Model 2 arises only for the highest-intensity genotypes: although *μ*(*i*) is small within the sublethal zone, it is nonzero, and because *d_p_* is small, this residual killing slightly lowers *P_2_*\*, so the decomposition assigns a share to the density channel. In Model 1, growth impairment has only a small effect on *P*\* because *d_p_*/*r*_eff_ is small. In contrast, in Model 2, *P*\* is independent of parasite growth rate in the sublethal zone, so growth impairment has no direct effect on parasite density. This insensitivity of *P*\* to growth impairment is part of the reason why the framework attributes most of the damage reduction to tolerance. The dominance of virulence reduction in Model 1 is conditional on *d_p_*/*r*_eff_ remaining small: it exceeds 90% while *d_p_*/*r*_eff_ ≲ 0.25 and declines as the ratio rises (see Supplementary Material); the ratio in the main analysis (0.054) lies far below this threshold. In Model 2, the same dominance is structural rather than parametric, since *P*\* does not depend on growth impairment. When the analysis was extended to include genotypes in the transition and killing zones, the framework’s difficulty persisted: transition-zone genotypes appeared maximally tolerant under linear damage, and killing-zone genotypes in Model 2 produced positive slopes under both damage functions (Supplementary Fig. S1).

**Table 2.** Fraction of total damage reduction of defence attributable to virulence reduction (Shapley decomposition) across host genotypes in the two model variants (Model 1, microparasite; Model 2, macroparasite) and damage functions (linear, sigmoidal).

| Genotype | M1 linear | M1 sigmoidal | M2 linear | M2 sigmoidal |
| --- | --- | --- | --- | --- |
| $\bar{i} = 0.05$ | 98.4% | 98.4% | 100.0% | 100.0% |
| $\bar{i} = 0.10$ | 98.3% | 98.3% | 99.7% | 99.7% |
| $\bar{i} = 0.15$ | 98.1% | 98.0% | 97.5% | 97.4% |
| $\bar{i} = 0.20$ | 97.4% | 97.2% | 90.5% | 90.0% |
| $\bar{i} = 0.25$ | 95.8% | 95.4% | 76.6% | 75.8% |

Across the full (*β_r_*, *β_α_*) parameter space, the dominance of virulence reduction was not specific to the choice *β*_r_ = *β*_α_ = 2.0. When *β_α_* = 0, immunity has no effect on virulence, and the virulence fraction is zero by definition (the dark stripe along the bottom edge of each panel of Fig. 2). As *β_α_* increases, virulence reduction rapidly dominates, its fraction exceeding 90% across most of the parameter space, depending on the model variant. In Model 2, this transition is sharp because parasite density is relatively insensitive to growth impairment: *P*\* is reduced only by immune killing, which is small in the sublethal zone, and is unaffected by growth impairment, so even a small value of *β_α_* dominates the damage reduction. In Model 1, the transition is more gradual because reducing the growth rate also slightly reduces *P*\*, so density reduction contributes even when *β_α_* is near zero.

**Figure 2.**
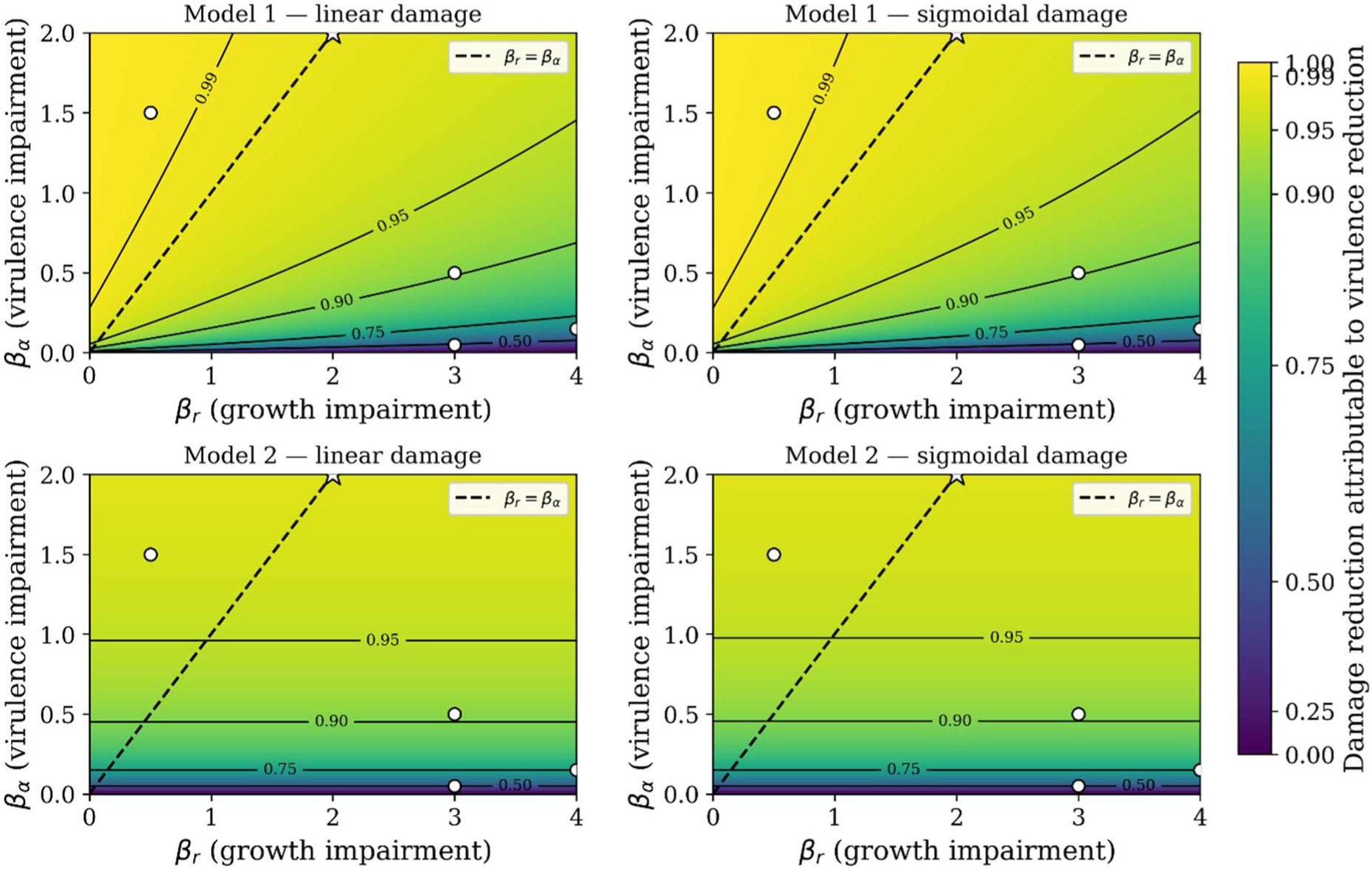
Fraction of total damage reduction attributable to virulence reduction (Shapley decomposition) across the (*β_r_*, *β_α_*) parameter space for Model 1 (microparasite, top) and Model 2 (macroparasite, bottom) under linear (left) and sigmoidal (right) damage functions. Star: *β_r_* = *β_α_* = 2.0 (main reaction norm analysis in Fig. 1). Open circles: four additional parameter combinations in which reaction norms were examined in detail (Supplementary Figs S2–S5).

The choice of damage function had virtually no effect on the decomposition: the linear and sigmoidal panels are nearly identical within each model variant (Fig. 2). This contrasts with Fig. 1, where the damage function reversed the slope rankings across genotypes. The damage function determines which genotype appears most tolerant (Fig. 1) but does not change the underlying decomposition as virulence reduction dominates the damage reduction regardless of whether damage is linear or sigmoidal (Fig. 2). Supplementary Figures S2–S5 confirm that the slope reversal shown in Figure 1 persists across four additional (*β_r_*, *β_α_*) combinations spanning different regions of the parameter space.

## Discussion

The central result of this study is that the reaction-norm framework cannot identify the mechanism underlying a shallow slope of host fitness on parasite burden. A host genotype whose immune system impairs parasites sublethally, thereby reducing per-parasite virulence without killing them, generates the same regression pattern that is classically interpreted as tolerance. More strikingly, the direction of the apparent tolerance gradient reverses between damage functions: under linear damage, the genotype with the strongest sublethal resistance appears most tolerant, whereas under sigmoidal damage, the genotype with the weakest resistance appears most tolerant. The result holds across both model variants (micro- and macroparasites), and the reversal of slope rankings persists across the examined (*β_r_*, *β_α_*) combinations. Virulence reduction accounts for more than 90% of the damage reduction across most of the parameter space, falling below that level where growth impairment is strong and virulence impairment is weak.

The structural reason is straightforward. Equilibrium parasite density is insensitive to sublethal resistance in both model variants. In Model 1, the ratio *d_p_*/*r*_eff_ is small, so *P*\* barely changes even when immune impairment substantially reduces the growth rate. In Model 2, *P*\* is entirely independent of growth rate in the sublethal zone because parasite density is set by the balance between environmental establishment and baseline mortality, neither of which depends on the parasite’s exploitation capacity. Because *P*\* barely changes across genotypes, most of the fitness variation appears as differences in *W* at roughly constant parasite load, which, by the operational definition of the framework, is tolerance. The same logic yields a more extreme outcome regarding immune costs (see Supplementary material). Near the clearance threshold, where parasite-induced damage is minimal regardless of load, fitness variation is dominated by immune costs. Thus, the reaction-norm slope can even turn positive. This pattern is interpretable as neither resistance nor tolerance, and further indicates that the reaction norm slope reflects whatever dominates fitness variation. Fitting more flexible, nonlinear tolerance curves does not, on its own, resolve this problem. Such curves decompose the fitness–load relationship into additional components (Louie et al., 2016; Gupta & Vale, 2017), but whether the regression is linear or nonlinear, it still cannot distinguish the mechanism underlying the observed relationship. The shape of the damage function, which is typically unknown, illustrates a separate difficulty: it determines the ranking of slopes across hosts with different levels of defence. Torres et al. (2016) and Lough et al. (2015) have argued that tracking host health and parasite load over time, rather than at a single time point, captures dynamics that static regressions miss. However, these approaches still compare host health against parasite load, and variation in health at a given load could reflect either host tissue protection or reduced per-parasite harm.

The resistance–tolerance dichotomy underpins a substantial body of evolutionary theory. Roy and Kirchner (2000) showed that resistance creates negative frequency-dependent selection, thereby maintaining polymorphism, whereas tolerance tends to fix in populations, thereby increasing disease prevalence. Restif and Koella (2004) demonstrated that mixed strategies can be evolutionarily stable when resistance and tolerance trade off. Miller et al. (2006) predicted that, where tolerance reduces infection-induced mortality, parasite virulence evolves upward in tolerant host populations, and Best et al. (2008) identified conditions under which genetic variation in tolerance is maintained. More recently, Singh and Best (2021) showed that, under a resistance–tolerance trade-off, faster host recovery and sterilising infections favour tolerance, whereas resistance is favoured otherwise, while Vitale and Best (2019) showed that reducing tolerance can paradoxically drive parasite extinction. These predictions all depend on resistance and tolerance being distinct, independently varying traits. Our results show that a single immune mechanism can generate both patterns simultaneously: impairment of parasite exploitation reduces per-parasite virulence (which the framework reads as tolerance) and, in Model 1, modestly reduces density (which the framework reads as resistance). If empirical estimates of ‘genetic variation in tolerance’ reflect sublethal resistance rather than parasite-independent tissue protection, then the evolutionary predictions derived from the dichotomy, including the expectation that tolerance increases prevalence (Boots, 2008) and does not provoke counter-adaptation (Råberg et al., 2009), may not apply in those systems.

If the reaction-norm framework alone cannot distinguish mechanisms, progress requires independent evidence that a given tolerance phenotype operates without affecting the parasite. Such mechanisms exist, though unambiguous examples remain scarce. Renal heme detoxification through ferritin-H and NRF2 limits kidney damage during malaria independently of pathogen burden (Ramos et al., 2019), and Soares et al. (2017) and Martins et al. (2019) review tissue-protective mechanisms, spanning metabolic adaptation, antioxidant defence, and tissue repair, that operate without targeting the parasite directly. However, many candidate tolerance mechanisms share molecular machinery with pathways that also impair parasites, making it difficult to conclusively establish parasite independence (McCarville & Ayres, 2018). The empirical challenge is to demonstrate that a candidate mechanism does not affect parasite condition, a negative result that requires measuring parasite fitness rather than just parasite density. Vale and Little (2012) provide a cautionary example. In *Daphnia* infected with *Pasteuria*, what appeared to be fecundity tolerance was in fact fecundity compensation. This host response resembled tolerance in the regression but operated through a distinct process. Integrating mechanistic immunology with reaction-norm analysis, rather than relying solely on regression, offers one way forward.

In systems where feasible, measuring parasite condition directly would provide the information the framework lacks. If parasites recovered from a ‘tolerant’ host are as fit as parasites from a naive host, the host’s defence is genuinely parasite-independent. If they are impaired, the host is exerting sublethal resistance that the framework cannot detect. Mackinnon and Read (2004) demonstrated the feasibility of this approach in rodent malaria, serially passaging *Plasmodium chabaudi* through semi-immune and naive mice and assaying parasite fitness in both host types. Birget et al. (2019) showed that multiplication rate, burst size, and cell-cycle duration are heritable, plastic parasite traits that respond measurably to host anaemia (red blood cell resource availability), whereas invasion preference for red blood cells of different ages showed neither genetic nor environmental variation. In helminth systems, per-capita egg output is a feasible proxy for worm condition, though it is confounded by density-dependent fecundity (Churcher et al., 2006). In *Drosophila*–pathogen systems, bacterial growth during the early phase of infection can be measured directly and predicts infection outcome (Duneau et al., 2017). None of these approaches is universally applicable, and in many natural systems—particularly wild vertebrate populations, where field studies of tolerance have been conducted (Hayward et al., 2014; Soler & Moller, 2023; Jackson et al., 2014)—direct measurement of parasite condition may be impractical. But they identify the type of additional data needed to mechanistically interpret a shallow slope, and because existing studies lack such data, their tolerance estimates cannot be assumed to reflect parasite-independent defence rather than undetected sublethal resistance.

Several limitations in our model should be noted. The model analyses equilibrium states, whereas real infections are dynamic processes in which resistance and tolerance may contribute at different phases (Lough et al., 2015; Duneau et al., 2025). We use a single scalar immune trait, collapsing the multidimensional reality of immune defence. Models with multiple interacting immune components could generate more complex patterns. The model does not consider coevolutionary dynamics. If parasites evolve in response to host immunity, the equilibrium parasite densities and virulence levels we analyse would shift, potentially altering the decomposition. *β_r_* has weak structural effects on *P*\* because baseline parasite mortality is small relative to the growth rate (Model 1) or because *P*\* is independent of the growth rate (Model 2). This limits the informativeness of the parameter space in the region where *β_r_* is high, and *β_α_* is low, though the biological relevance of that region is itself questionable. Finally, the sigmoidal damage function is one of many possible nonlinear forms; other shapes could produce different slope patterns, and the finding that linear and sigmoidal functions produce opposite tolerance rankings should be understood as an illustration of damage-function dependence, not a broad survey.

To conclude, the slope of a reaction norm does not identify the mechanism underlying a host’s response to infection. A shallow slope is consistent with genuine tissue-protection tolerance, sublethal immune resistance that impairs parasites without killing them, or any combination of the two. The reaction-norm framework provides a useful description of the statistical relationship between host fitness and parasite load, but it cannot, without additional information, distinguish the mechanisms that generate this relationship. Moving forward requires two things: (1) identifying tolerance mechanisms that are demonstrably parasite-independent, using the molecular and physiological tools now available (Soares et al., 2017; Martins et al., 2019), and (2) measuring parasite condition alongside host fitness and parasite load in systems where this is feasible. Until both are in place, claims of genetic variation in tolerance based on reaction-norm slopes alone should be interpreted with caution.

## Supporting information

Supplementary analyses

## Acknowledgments

We thank K. King for helpful discussions.

