## Supplementary analyses for "Sublethal immune resistance to parasites generates reaction-norm patterns indistinguishable from tolerance"

**Supplementary Material for '*Sublethal immune resistance to parasites generates reaction-norm patterns indistinguishable from tolerance*'**

### Supplementary analyses

*1. Extension to the full range of immune intensities*

The main analysis is restricted to the sublethal zone, where killing is negligible, and the main immune effect is impairment of parasite condition. This provides a clean demonstration that sublethal resistance alone generates patterns classified as tolerance. However, natural populations include genotypes whose immune responses are strong enough to kill parasites. Here, we ask whether the framework's difficulty persists, diminishes, or worsens when the analysis includes such genotypes. We added two genotypes to the five used in the main analysis: $\bar{\imath} = 0.40$ (transition zone, where killing contributes substantially but does not yet dominate: $\mu = 0.12$) and $\bar{\imath} = 0.65$ (killing zone, above the half-maximal threshold $\theta\mu = 0.6$: $\mu = 0.93$). All other aspects of the simulation (three parasite clones, 20 replicates per cell, two damage functions) were identical to those in the main analysis.

Model 1 (microparasite): the killing-zone genotype ($\bar{\imath} = 0.65$) cleared all infections and was therefore dropped from the analysis (no infected individuals could be assigned a tolerance slope). Under linear damage, the transition-zone genotype ($\bar{\imath} = 0.40$) had the shallowest slope (−4.6), making it appear the most "tolerant" genotype in the dataset, although it induces substantial killing ($\mu = 0.12$; Fig. S1a). The shallow slope arises because per-parasite virulence is low: each additional parasite causes little damage, and fitness changes slowly with load. Under sigmoidal damage, the same genotype no longer appeared tolerant at all as its slope steepened to −20.4 (from −4.6 under linear damage), placing it among the least tolerant genotypes. As under sigmoidal damage in the main analysis, the apparent-tolerance ranking reverses relative to linear damage.

Model 2 (macroparasite): both additional genotypes remained in the analysis, because macroparasites continue to arrive from the environment even under strong killing. The killing-zone genotype ($\bar{\imath}$ = 0.65) had very low parasite density ($P^*$ = 0.8–3.7) and parasites retained only 27% of their baseline exploitation capacity. Under both damage functions, its reaction-norm slope was positive (Fig. S1c,d). The positive slopes arise because, at the very low parasite densities characteristic of the killing zone, parasite-induced damage is minimal regardless of load. Thus, the dominant source of fitness variation among individuals of this genotype is the immune cost: individuals with slightly higher $i$ (due to within-genotype noise) pay higher costs, creating a positive association between $P^*$ and $W$. This is a structural limitation of the framework when applied to hosts near the clearance threshold.

The extension to the full intensity range reveals two additional problems beyond those demonstrated in the main analysis. First, genotypes that clear infection disappear from the analysis (Model 1, killing zone), creating a survivorship bias: the framework can only evaluate hosts that fail to eliminate their parasites. Second, genotypes near the clearance threshold can produce positive slopes driven by immune defence costs rather than by any tolerance mechanism (Model 2, killing zone).

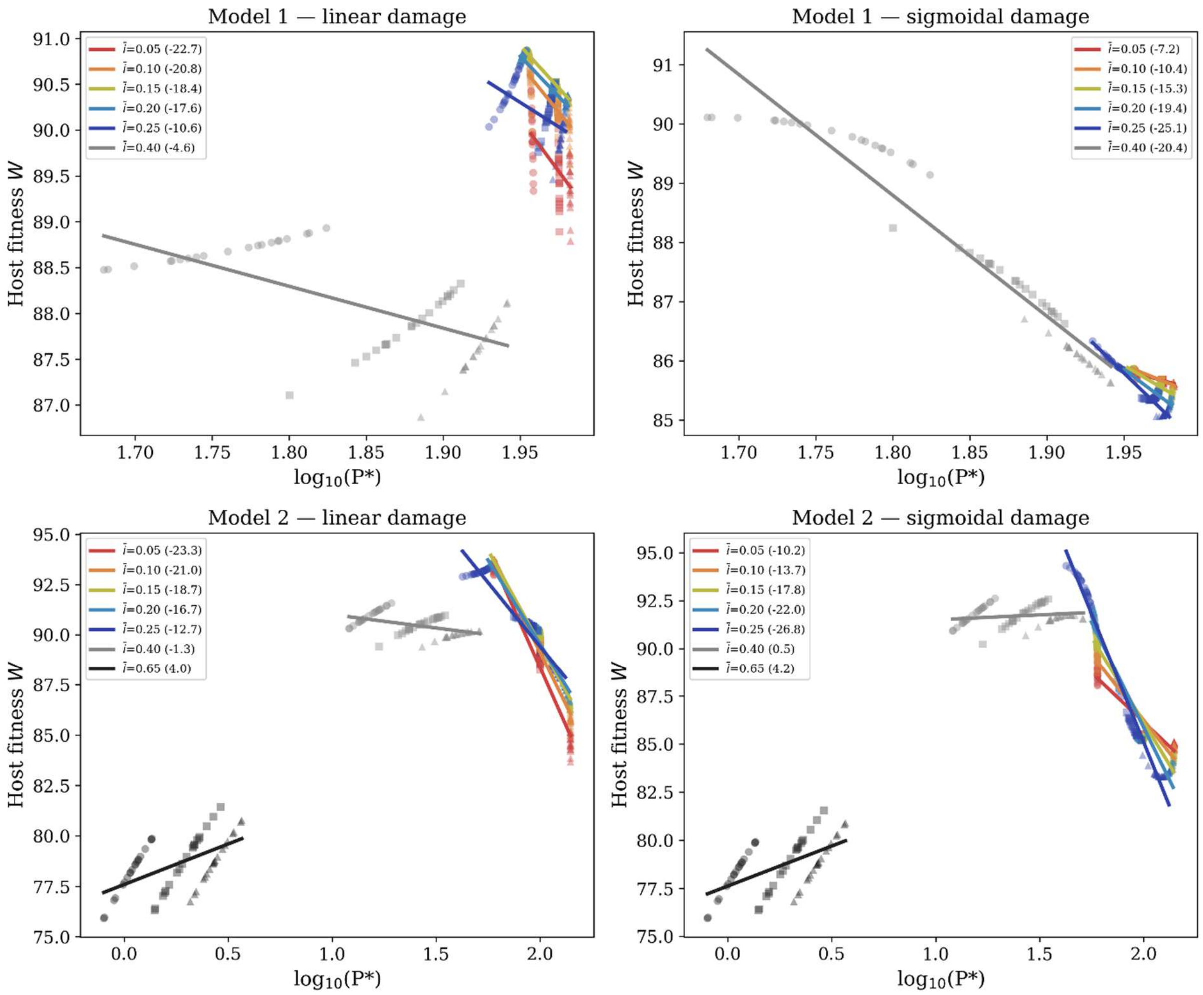


**Figure S1.** Reaction norms across the full range of immune intensities for Model 1 (microparasite, top) and Model 2 (macroparasite, bottom) under linear (left) and sigmoidal (right) damage functions. The five original sublethal genotypes are coloured as in the main figure. Grey = transition zone ($\bar{\imath}$ = 0.40), black = killing zone ($\bar{\imath}$ = 0.65). Markers indicate parasite clones with different reproductive rates: ○ slow, □ medium, △ fast.

*2. Reaction norms at alternative combinations of parameter values*

We repeated the reaction-norm analysis at four additional ($\beta_r$, $\beta_\alpha$) combinations spanning different regions of the parameter space (marked as open circles in Fig. 2). When virulence was more sensitive to immunity than growth ($\beta_r = 0.5$, $\beta_\alpha = 1.5$; Fig. S2), the slope reversal between damage functions was pronounced, and virulence reduction accounted for 70–100% of the damage reduction. Reversing the balance so that growth was more sensitive ($\beta_r = 3.0$, $\beta_\alpha = 0.5$; Fig. S3) reduced the virulence fraction to 45–100%, but the slope reversal persisted. Even near the bottom edge of the parameter space, where virulence impairment was minimal ($\beta_r = 3.0$, $\beta_\alpha = 0.05$; Fig. S4), the slope reversal was present, although virulence reduction now contributed only about half the damage reduction in Model 1 (31–52%) and as little as 8% in Model 2. A fourth combination with strong growth-rate suppression and modest virulence impairment ($\beta_r = 4.0$, $\beta_\alpha = 0.15$; Fig. S5) produced similar results: virulence reduction contributed 49–70% in Model 1 and 20–100% in Model 2, and the slope reversal held. In all four cases and under both damage functions, the ranking of genotypes reversed between linear and sigmoidal damage, confirming that the result demonstrated in Figure 1 is not specific to the choice $\beta_r = \beta_\alpha$.

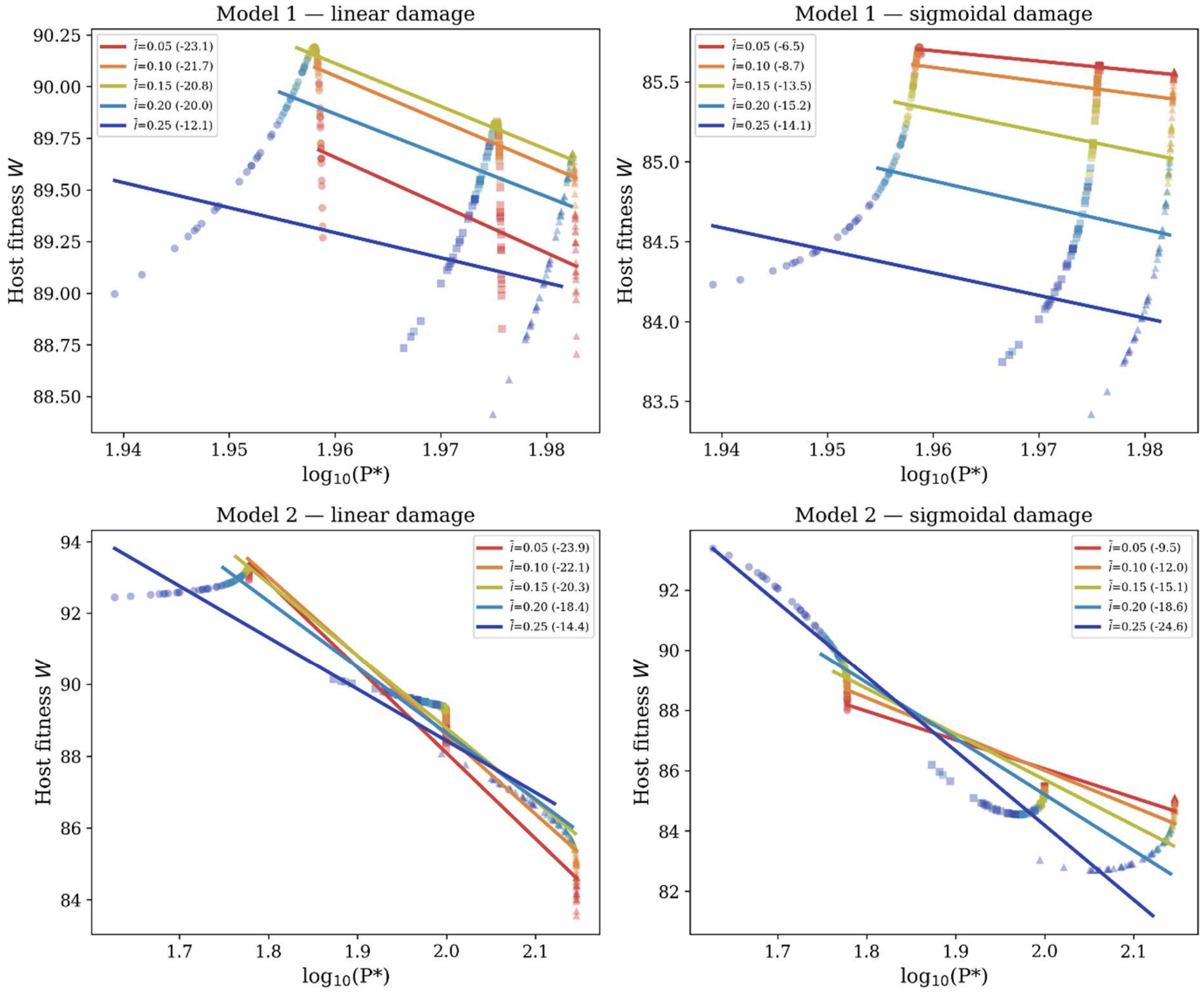


**Figure S2.** Reaction norms with $\beta_r = 0.5$, $\beta_\alpha = 1.5$ (above the diagonal in Fig. 2; virulence is more sensitive to immunity than growth). Markers indicate parasite clones with different reproductive rates: ○ slow, □ medium, △ fast.

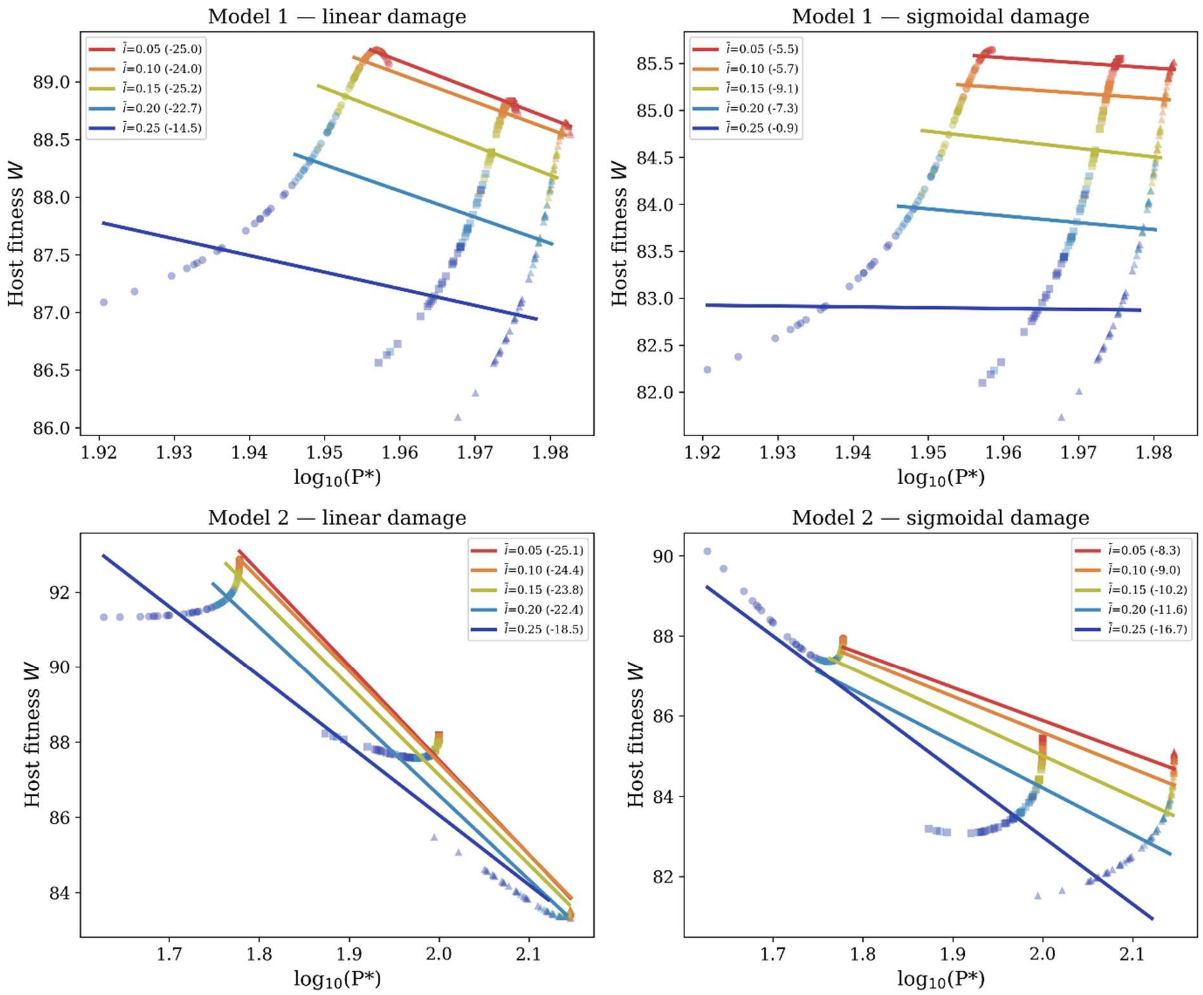


**Figure S3.** Reaction norms with $\beta_r$ = 3.0, $\beta_\alpha$ = 0.5 (below the diagonal in Fig. 2; growth is more sensitive to immunity than virulence). Markers indicate parasite clones with different reproductive rates: ○ slow, □ medium, △ fast.

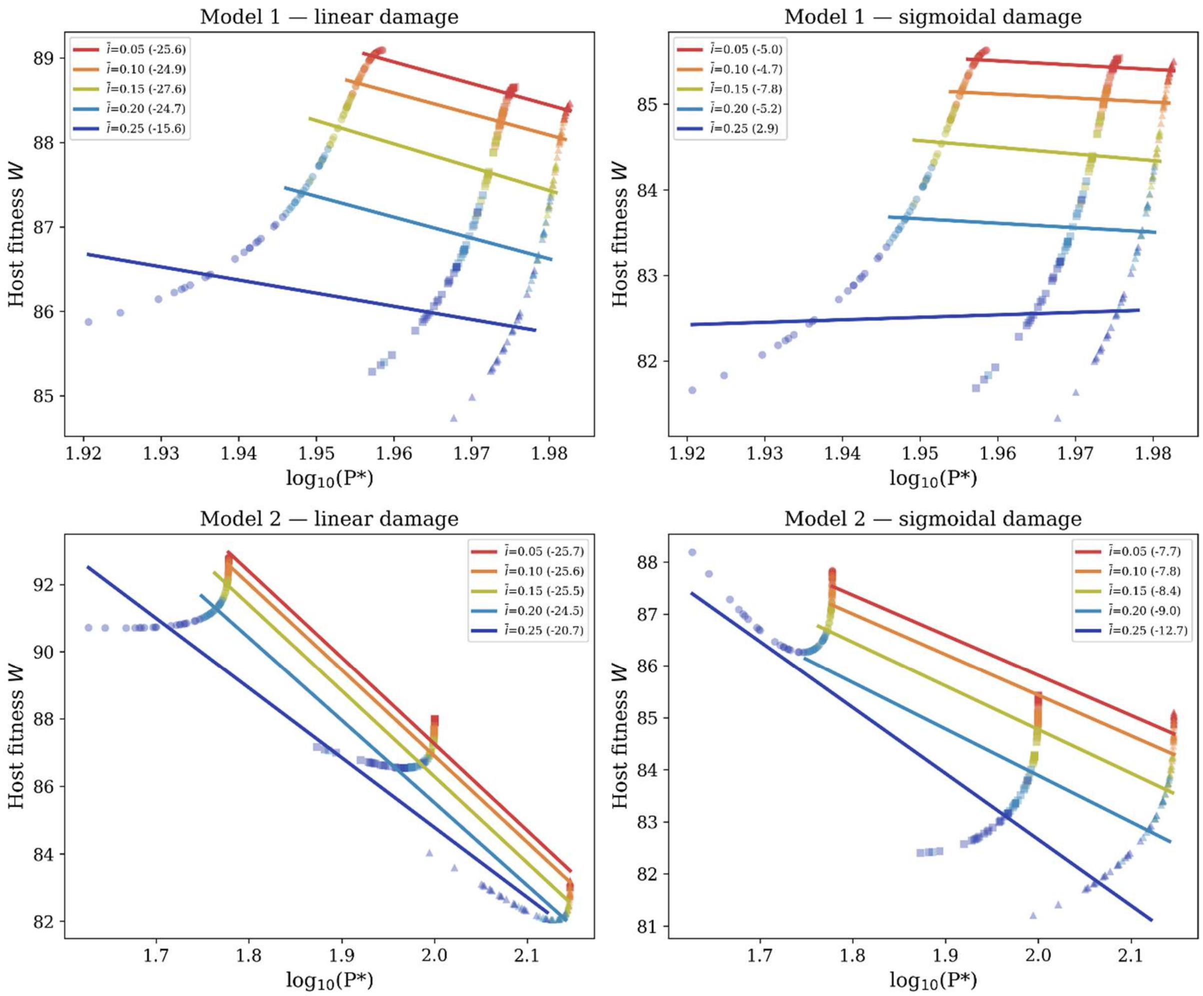


**Figure S4.** Reaction norms with $\beta_r = 3.0$, $\beta_\alpha = 0.05$ (near the bottom edge of Fig. 2; strong growth impairment, minimal virulence impairment). Markers indicate parasite clones with different reproductive rates: ○ slow, □ medium, △ fast.

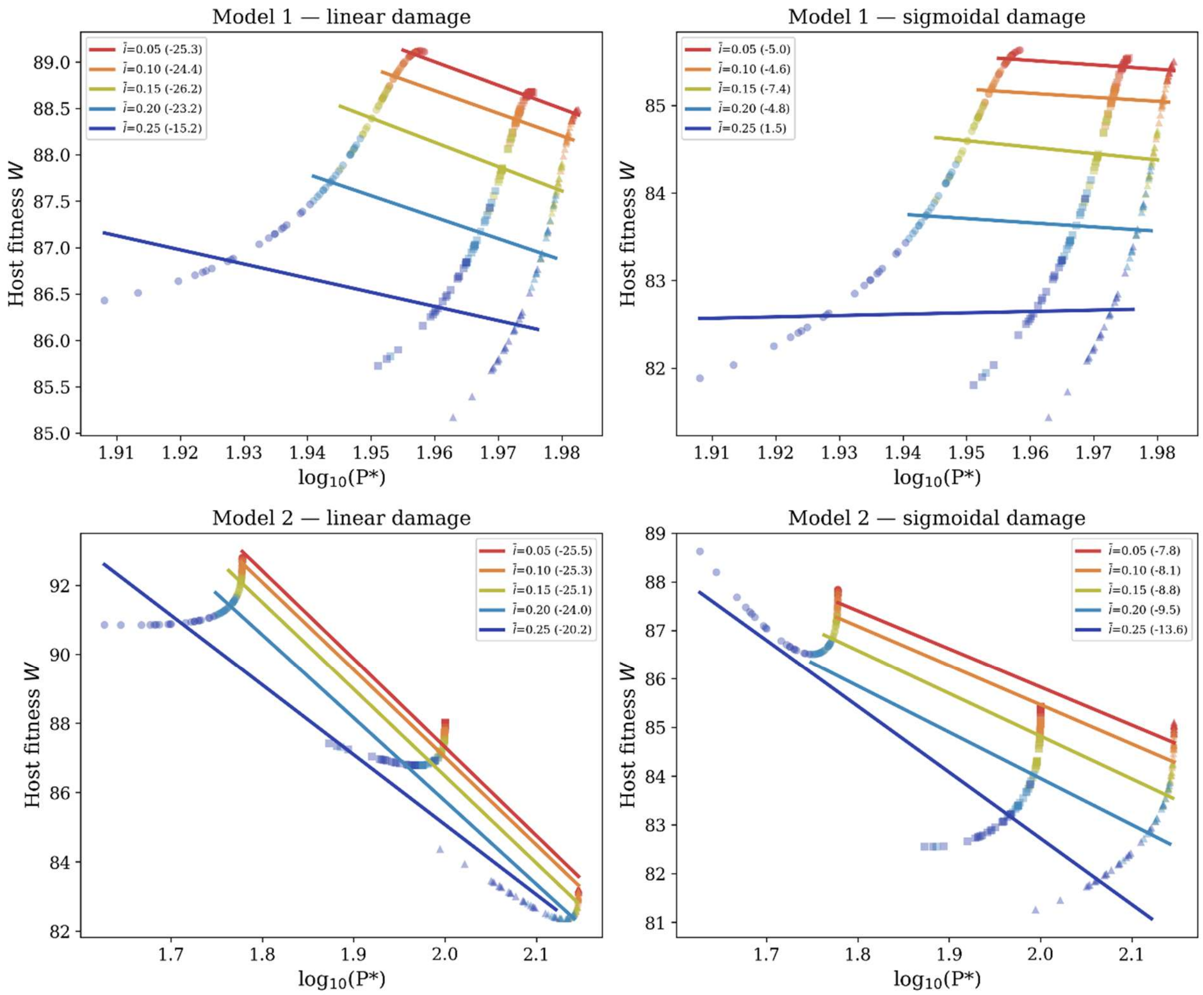


**Figure S5.** Reaction norms with $\beta_r = 4.0$, $\beta_\alpha = 0.15$ (lower right of Fig. 2; strong growth impairment, modest virulence impairment). Markers indicate parasite clones with different reproductive rates: ○ slow, □ medium, △ fast.

*3. Sensitivity of the decomposition to the ratio $d_p/r_{eff}$*

The decomposition in the main analysis assumes that equilibrium parasite density is nearly insensitive to growth impairment. In Model 1, this holds only while baseline parasite mortality is small relative to the effective growth rate, $d_p/r_{\mathrm{eff}} \ll 1$. To test how far this assumption can be relaxed, we recomputed the Model 1 virulence-reduction fraction ($\bar{\iota} = 0.15$, $\beta_r = \beta_\alpha = 2.0$, averaged over clones) while varying $d_p$, and expressed the result against the baseline ratio $d_p/r_{\mathrm{eff}}$ (Fig. S6). Virulence reduction accounted for more than 90% of the damage reduction for $d_p/r_{\mathrm{eff}}$ up to about 0.25 and declined smoothly as the ratio rose, reaching roughly 60% by $d_p/r_{\mathrm{eff}} = 0.55$; the value used in the main analysis ($d_p/r_{\mathrm{eff}} = 0.054$) therefore lies well within the regime where virulence reduction dominates, and the result was nearly identical under linear and sigmoidal damage. This dependence follows from $\partial P^*/\partial r_{\mathrm{eff}} \propto d_p/r_{\mathrm{eff}}^2$: when $d_p/r_{\mathrm{eff}}$ is small, growth impairment barely moves $P^*$, so the density channel is negligible and almost all of the damage reduction is assigned to virulence reduction. As the ratio grows, growth impairment lowers $P^*$ appreciably, and the density channel begins to contribute. Beyond $d_p/r_{\mathrm{eff}} \approx 0.6$, the slowest clone reaches $P^* = 0$ at baseline, so the parasite cannot persist, and the decomposition is undefined. In Model 2, no analogous analysis is required as $P^*$ is set by the balance between environmental establishment and baseline mortality and does not depend on growth impairment. The dominance of virulence reduction in Model 2 is structural rather than parametric.

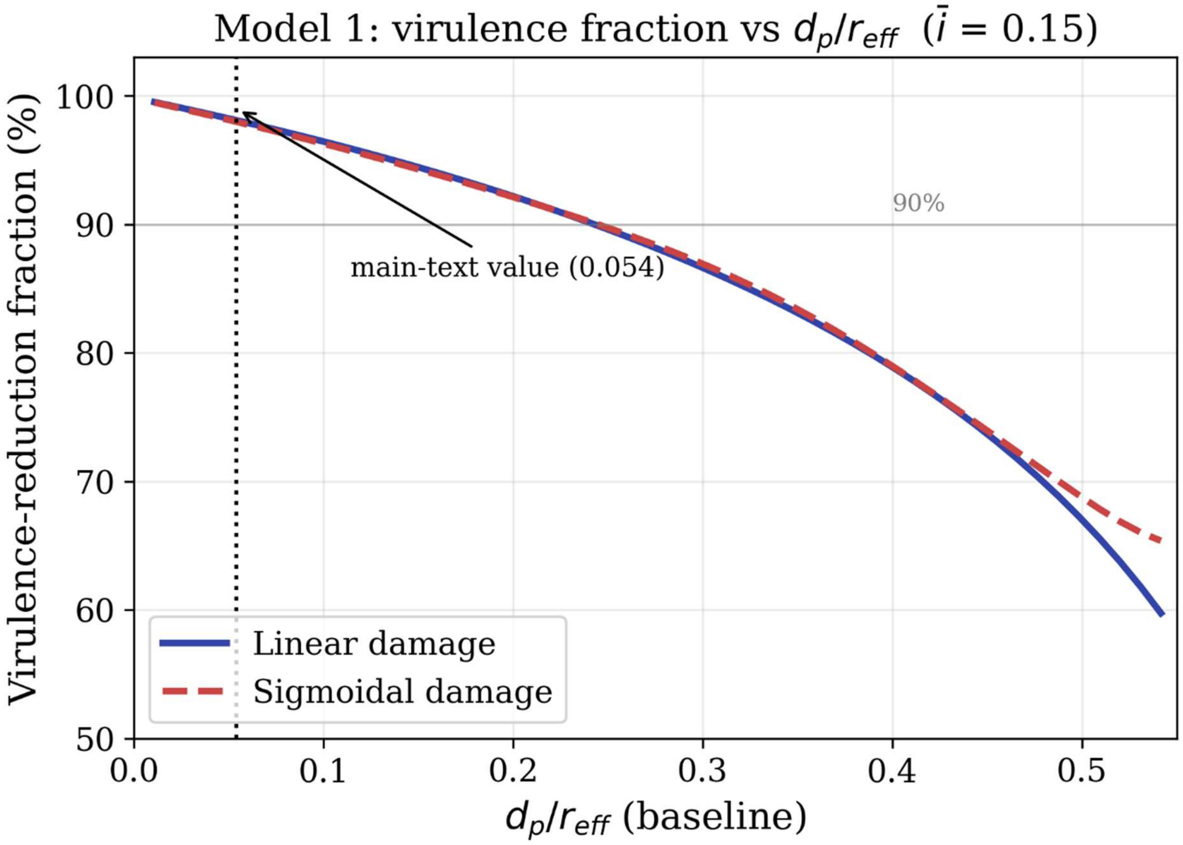


**Figure S6.** Sensitivity of the Model 1 decomposition to the ratio $d_p/r_{\mathrm{eff}}$. The virulence-reduction fraction (Shapley decomposition; $\bar{\imath} = 0.15$, $\beta_r = \beta_\alpha = 2.0$, averaged over clones) as a function of the baseline ratio $d_p/r_{\mathrm{eff}}$ for linear (solid) and sigmoidal (dashed) damage. The dotted line marks the value used in the main analysis (0.054).
